# Gold Nanoparticles interacting with Synthetic Lipid Rafts: an AFM investigation

**DOI:** 10.1101/2020.05.03.075101

**Authors:** Andrea Ridolfi, Lucrezia Caselli, Costanza Montis, Gaetano Mangiapia, Debora Berti, Marco Brucale, Francesco Valle

## Abstract

Inorganic nanoparticles (NPs) represent promising examples of engineered nanomaterials, providing interesting biomedical solutions in several fields, like therapeutics and diagnostics. Despite the extensive number of investigations motivated by their remarkable potential for nanomedicinal applications, the interactions of NPs with biological interfaces are still poorly understood. The effect of NPs on living organisms is mediated by biological barriers, such as the cell plasma membrane, whose lateral heterogeneity is thought to play a prominent role in NPs adsorption and uptake pathways. In particular, biological membranes feature the presence of rafts, i.e. segregated lipid micro and/or nano-domains in the so-called liquid ordered phase (L_o_), immiscible with the surrounding liquid disordered phase (L_d_). Rafts are involved in various biological functions and act as sites for the selective adsorption of materials on the membrane. Indeed, the thickness mismatch present along their boundaries generates energetically favorable conditions for the adsorption of NPs. Despite its clear implications in NPs internalization processes and cytotoxicity, a direct proof of the selective adsorption of NPs along the rafts’ boundaries is still missing to date. Here we use multicomponent Supported Lipid Bilayers (SLBs) as reliable synthetic models, reproducing the nanometric lateral heterogeneity of cell membranes. After being characterized by Atomic Force Microscopy (AFM) and Neutron Reflectivity (NR), multi-domain SLBs are challenged by prototypical inorganic nanoparticles, i.e. citrated gold nanoparticles (AuNPs), under simplified and highly controlled conditions. By exploiting AFM, we demonstrate that AuNPs preferentially target lipid phase boundaries as adsorption sites. The herein reported study consolidates and extends the fundamental knowledge on NPs-membrane interactions, which constitute a key aspect to consider when designing NPs-related biomedical applications.

## INTRODUCTION

Despite the impressive technological advancement in the design of “smart” inorganic nanoparticles (NPs), their impact on biological systems and related toxicity are still poorly understood^1,2^, limiting their effective clinical translation. The interaction of engineered nanomaterials, either intentionally or inadvertently released into the environment, with living organisms is mediated by biological barriers, such as cell plasma membranes, which primarily determine NPs biological fate and cytotoxicity^3^. Therefore, the interaction of NPs with biological interfaces is a key research topic, aiming at the safe use of nanotechnology and maximization of its potential in therapeutics and diagnostics^4,5^.

In this framework, lipid-based synthetic model membranes are useful platforms to mimic biological interfaces under simplified conditions, allowing for the identification of key determinants regulating nano-bio interactions^6–8^. Supported Lipid Bilayers (SLBs) are often used as 2D biomembrane models^9,10^, enabling to precisely tune their physicochemical properties and avoiding the complications related to the 3D nature of biological membranes. They also represent versatile and promising platforms for the development of biosensors^11^ and technological assays for biological applications^12^.

In addition, multicomponent SLBs models allow studying the lateral compositional heterogeneity that characterizes most biological membranes. The presence of discrete lipid domains in natural membranes is thought to have a profound impact on their biological function, as exemplified by cells^13^. A specific case of lateral organization is represented by lipid rafts, defined as micro and/or nano-domains, enriched in lipids such as cholesterol, sphingomyelin, saturated glycerophospholipids and glycosphingolipids: these lipids segregate in the so-called liquid-ordered phase (L_o_), which is immiscible with the surrounding liquid-crystalline (disordered, L_d_) phase^14^. This phase heterogeneity induces a thickness mismatch between neighboring domains and the consequent, ergonically unfavorable, exposure of hydrocarbon regions to water, which results in an energetic cost, due to interfacial energy^15^. Rafts are thought to participate in the formation and targeting of nano-sized biogenic lipid vesicles (e.g. Extracellular Vesicles, EVs)^16^. They are also actively involved in multiple membrane processes, e.g. they act as structural platforms for organizing protein machinery^17^, they can preferentially associate with specific membrane proteins^18^ and represent centers for the assembly of signaling molecules. From a mechanical point of view, the presence of phase boundaries and, hence, bilayers thickness mismatches, generates deformations and increases membrane permeability^19–21^. All these structural perturbations promote the selective adsorption of materials on the membrane; indeed, as pointed out by Hamada et al.^22^, lateral heterogeneity, promoted by the presence of micro-sized lipid rafts, regulates the adsorption of nano/microparticles, with the larger ones preferring the L_d_ phases and the smaller ones being localized in the L_o_ phases of cell-sized lipid vesicles. These selective NPs adsorption pathways are also present in the case of nano-sized lipid segregated domains and can be studied exploiting liposomes with tunable rafts size^15^. However, investigating the interaction of NPs with nanometric lipid rafts remains a major challenge, mainly hindered by the small size of the segregated domains, which makes standard optical techniques not suitable for the task. Recent studies demonstrated that gold nanoparticles (AuNPs) adsorb more strongly to phase-separated multicomponent lipid bilayers; in particular, they are believed to preferentially target the phase boundary regions, due to their intrinsic negative curvature^23,24^.

To the best of our knowledge, this behavior has only been investigated by computational studies^23^ and experiments involving Quartz Crystal Microbalance (QCM)^24^, which provide important but indirect evidences. In summary, the preferential adsorption of AuNPs along the boundaries of nano sized lipid domains has never been directly observed.

To fill this gap, we exploit Atomic Force Microscopy (AFM), to directly visualize the preferential adsorption of AuNPs on the phase boundaries of multicomponent SLBs, presenting both an L_d_ and an L_o_ phase and previously characterized by Neutron Reflectivity (NR). The L_d_ phase is made of 1,2-dioleoyl-sn-glycero-3-phosphocholine (DOPC) with two unsaturated hydrocarbon chains that hinder molecular packing, while the L_o_ phase is composed of 1,2-dipalmitoyl-sn-glycero-3-phosphocholine (DSPC) lipids and cholesterol molecules which occupy the free volume between the lipid acyl chains^13,25^. The quantitative localization and morphometry of AuNPs adsorbed on the SLB reveal important information regarding their interaction with the lipid matrix. The study corroborates the already theorized differential NPs-lipid interaction taking place at the phase boundaries of lipid rafts. The presented results could help the development of future NPs-based applications that involve their adsorption on membranes characterized by nanoscale phase segregations.

## MATERIALS AND METHODS

### Materials

Tetrachloroauric (III) acid (≥ 99.9%), trisodium citrate dihydrate (≥ 99.9%), methanol (99.8%), CHCl_3_ (≥ 99.9%), NaCl (≥99.5%) and CaCl_2_ (99.999%) were provided by Sigma-Aldrich (St. Louis, MO). The same for 1,2-dioleoyl-sn-glycero-3-phosphocholine (DOPC) (≥ 98.0%), cholesterol (≥99.5%) and 1,2-distearoyl-sn-glycero-3-phosphocholine (DSPC) (≥ 98.0%). All chemicals were used as received. Milli-Q grade water was used in all preparations.

### AuNP preparation

Anionic gold nanospheres of 16 nm in size were synthesized according to the Turkevich-Frens method^26,27^. Briefly, 20 ml of a 1 mM HAuCl_4_ aqueous solution were brought to boiling temperature under constant and vigorous magnetic stirring. 2 ml of 1% citric acid solution were then added and the solution was further boiled for 20 minutes, until it acquired a deep red color. The nanoparticles solution dispersion was then slowly cooled down to room temperature.

#### Vesicle preparation and SLB formation for Neutron Reflectivity measurements

##### Vesicle preparation

The proper amount of a DOPC/DSPC/cholesterol mixture (39/39/22 mol%) was dissolved in chloroform and a lipid film was obtained by evaporating the solvent under a stream of nitrogen and overnight vacuum drying. The film was then swollen and suspended in warm (50 °C) 100 mM NaCl water solution by vigorous vortex mixing, in order to obtain a final 0.5 mg/ml lipid concentration. The resultant multilamellar vesicles (MLVs) were tip sonicated with a Digital Sonifier Model 450 (Branson), provided with a Horn Tip (diameter 25.4 mm), in an intermittent-pulse mode (5 seconds), with a power of 400 W (amplitude 50%), for 15 minutes to obtain a homogeneous dispersion of unilamellar vesicles (ULVs).

##### Surface cleaning procedure

DOPC/DSPC/cholesterol single lipid bilayers were formed on 50 × 80 × 15 mm^3^ Silicon mirrors (Andrea Holm GmbH, Tann, Germany; roughness ≤ 5 Å). Substrates were preliminary rinsed in either ultrapure water and ethanol, in order to remove organic residues. After that, they were bath sonicated treated for 30 minutes in ethanol with a Bandelin DL 102 3L bath sonicator, followed by other 30 minutes in ultrapure water (Millipore Simplicity UV). The surfaces were then cleaned with a Novascan PSD-UV8 UV/ozone plasma for 30 min and rinsed in ultrapure water. Finally, they were dried with nitrogen gas and stored in ultrapure water, ready for the deposition.

##### Vesicle fusion and SLB formation

CaCl_2_ was added to the vesicle dispersion, reaching a final concentration of 10 mM, just before the injection in the NR measuring cell. This was performed in order to promote their adhesion to the support and their subsequent disruption. Vesicles were left incubating for 30 minutes; then, the saline buffer was switched to D_2_O to promote the vesicle disruption and SLB formation. The use of D_2_O instead of ultrapure water ensures a better resolution of the lipid structures for the NR measurements.

#### Vesicle preparation and SLB formation for AFM measurements

##### Vesicle preparation

The proper amount of a DOPC/DSPC/cholesterol mixture (39/39/22 mol%) was dissolved in chloroform and a lipid film was obtained by evaporating the solvent under a stream of nitrogen and overnight vacuum drying. The film was then swollen and suspended in warm (50 °C) ultrapure water solution by vigorous vortex mixing, in order to obtain a final 0.5 mg/ml lipid concentration. The resultant multilamellar vesicles in water were subjected to 10 freeze-thaw cycles and extruded 10 times through two stacked polycarbonate membranes with 100 nm pore size at room temperature, to obtain unilamellar vesicles with narrow and reproducible size distribution. The filtration was performed with the Extruder (Lipex Biomembranes, Vancouver (Canada)) through Nuclepore membranes.

##### Surface cleaning procedure

All reagents were purchased from Sigma-Aldrich Inc (www.sigmaaldrich.com). DOPC/DSPC/Chol supported lipid bilayers were formed on microscopy borosilicate glass coverslips (Menzel Gläser). Glass slides were first immersed in a 3:1 mixture of 96% H_2_SO_4_ and 50% aqueous H_2_O_2_ (‘oxidizing piranha’) solution for 2h in order to remove any organic residue present on their surface. Then, the slides were cleaned in a sonicator bath (Elmasonic Elma S30H) for 30 minutes in acetone, followed by 30 minutes in isopropanol and 30 minutes in ultrapure water (Millipore Simplicity UV). Glass slides were then cleaned with air plasma for 15 minutes (Air plasma cleaner PELCO easiGlow) and incubated in ultrapure water for 10 minutes in order to maximize the number of reactive silanols present on the surface. Finally, they were dried with nitrogen gas and stick to a magnetic disk, ready for the lipid solution deposition.

##### Vesicle fusion and SLB formation

A 100 μl droplet of buffer solution was firstly spotted on the SiO_2_ slide. The buffer solution consisted of CaCl_2_ 200 mM diluted 1:10 in KCl 100 mM. A 10 μl droplet containing the lipid mixture was then added to the buffer droplet and left incubating at room temperature for 30 minutes in order to promote the vesicle adsorption on the surface. After that, the droplet was removed and replaced by a 100 μl droplet of ultrapure water which was then left incubating for other 15 minutes. AuNPs deposition on the SLB was obtained by adding 5 μl of a 7.8 nM AuNPs dispersion to the ultrapure water droplet and leaving it to incubate for 10 minutes. After the system equilibrated, the large droplet was gently removed and the slide was inserted in the AFM fluid cell for the measurements.

##### Neutron Reflectivity measurements

NR measurements were conducted at the REFSANS Horizontal TOF reflectometer of the Helmholtz-Zentrum Geesthacht located at the Heinz Maier-Leibnitz Zentrum in Garching, Germany^28,29^. Neutrons in the wavelength range 3.0–21.0 Å were used to carry out the measurements. Two incident angles, namely 0.60° and 3.00°, allowed collecting data in the range 0.007 ≤ Q/Å^−1^ ≤ 0.22. The arrival times and positions of scattered neutrons were detected on a Denex 2D 500 × 700 mm^2^ multiwire ^3^He detector (pixel size 2.1×2.9 mm^2^, efficiency 80% at 7 Å, gamma sensitivity < 10^−6^) positioned at 4.5 m from the sample. The detector was installed in a liftable vacuum tube in order to reach exit angles up to 5.2° at the maximum elongation. In order to receive sufficient statistics, a counting time of about 4 hours for the measurement was chosen. The software MOTOFIT^30^ was employed for the analysis of the NR curve. Details on data analysis are reported in the SI.

#### AFM measurements

##### AFM setup

All AFM experiments were performed at room temperature on a Bruker Multimode 8 (equipped with Nanoscope V electronics, a sealed fluid cell and a type JV piezoelectric scanner) using Bruker SNL-A probes (triangular cantilever, nominal tip curvature radius 2-12 nm, nominal elastic constant 0.35 N/m) calibrated with the thermal noise method^31^. The AFM fluid cell was filled with a saline buffer solution, consisting of KCl 100 mM, which has the main effect of reducing the Debye length that characterizes the Electrical Double Layer (EDL) interaction region between AFM tip and SLB^32^. In this way, better image resolution can be achieved.

##### AFM Imaging

Imaging was performed in PeakForce mode. In order to minimize deformations or rupture events induced by the scanning probe, the applied force setpoint was kept under 200 pN range. Feedback gain was set on values high enough to obtain optimal image quality but low enough to prevent the introduction of noise signals that would otherwise interfere with the resolution of the different lipid domains (lipid phases have a height difference of ~1 nm).

The average height value of all bare substrate zones was taken as the baseline zero height reference. Image background subtraction was performed using Gwyddion 2.53.16^33^.

## RESULTS AND DISCUSSION

### Formation of Supported Lipid Bilayers containing lipid rafts

The formation of a continuous planar bilayer (DOPC/DSPC/cholesterol (39/39/22 mol%)), covering the vast majority of the supporting surface, was achieved through vesicle fusion and characterized by NR. Briefly, as described in the Materials and Methods section, liposomes in a saline buffer were additioned with a low amount of CaCl_2_, injected within the measurement chamber and left adsorbing on the support (a clean Si crystal). The presence of Ca^2+^ ions in solution promotes the crowding of vesicles on the surface by reducing the repulsive interactions between liposomes with surface charge. As reported by Richter et al.^9^, when a critical vesicle coverage is reached, the stress on the vesicles becomes sufficient to induce their rupture; in our case the phenomenon was also favored by the additional osmotic shock, coming from the replacement of the saline buffer with ultrapure water. The edges of the newly-formed SLB are energetically unfavorable and cause the rupture of other surface-bound vesicles. If the density of adsorbed vesicles is sufficiently high, these cascade phenomena can lead to the complete surface coverage.

Neutron Reflectivity (NR) was applied to probe the effective formation of the multicomponent SLB and its structure along the normal to the SLB plane. Figure 1a shows a representative NR profile measured for the SLB in D_2_O (green circles), together with the fitting curve (red continuous line). The curve was analyzed with MOTOFIT and, consistently with the literature^34–36^, it was possible to model the profile of the SLB as a stack of five layers (see scheme in Figure 1b): the silicon oxide layer, a layer of solvent (D_2_O), a layer for the polar headgroups in contact with the support (inner heads), a layer for the lipid chains (chains) and, finally, a layer for the polar headgroups in contact with the solvent (outer heads). Each layer is characterized by a defined contrast (the scattering length density, SLD), thickness (d), roughness (ρ) and hydration (solvent %). The curve fitting results are reported in Table 1. The overall thickness of the bilayer is ~ 5 nm (given by the sum of the thickness values related to the inner and outer heads, plus the lipid chains). The negligible hydration (0.1 %) of the lipid chains layer indicates that the surface was almost completely covered by a homogeneous lipid bilayer. The analysis of the experimental data allowed reconstructing the entire profile of the SLB along the normal to the surface (see Figure 1b).

**Figure 1.**
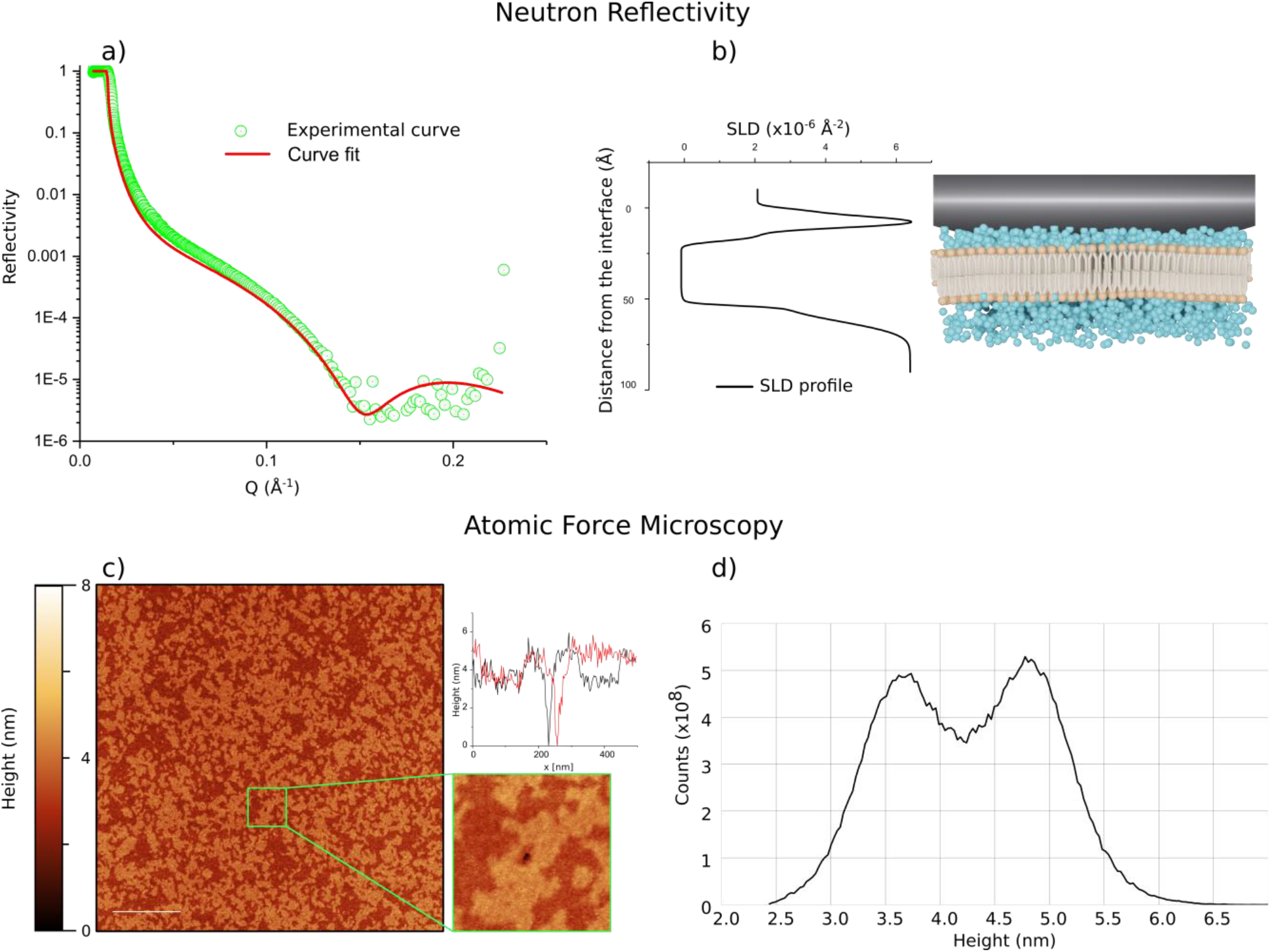
Characterization of the multicomponent SLB formed from DOPC/DSPC/cholesterol (39/39/22 mol%) liposomes by vesicle fusion. a) Neutron Reflectivity profile (green circles) and best fit (continuous red line) corresponding to the SLB in D_2_O; from the fitting analysis the average bilayer thickness is ~ 5 nm. b) Scattering length density (SLD) profile, describing variations of the SLD along the direction perpendicular to the bilayer. c) Representative AFM topography of the SLB. The bilayer uniformly covers the surface, displaying both the L_o_ (brighter thicker regions) and Ld phases (darker thinner regions) as segregated domains. The reported scalebar is 1 μm. The 500×500 nm micrograph (bottom inset) displays the small hole in the bilayer that allowed flattening the image with respect to the SiO_2_ surface. Two perpendicular height profiles were traced, horizontally and vertically, across the whole image (top inset); the profiles confirm the presence of the two distinct lipid phases covering the surface. d) Height distribution obtained from the AFM image; the two distinct peaks, centered at h_d_ = 3.7 nm and h_o_ = 4.7 nm, describe the different heights that characterize the L_d_ and L_o_ phase, respectively.

**Table 1.** Curve fitting results of NR data, obtained with MOTOFIT. The reported fitting parameters are referred to the three layers composing the bilayer (Inner Heads, referred to the layer of polar headgroups in contact with the support, Lipid Chains, referred to the hydrophobic region of the SLB, Outer Heads, referred to the layer of polar headgroups in contact with the solvent (i.e., D_2_O)). For each layer four parameters are reported: d(Å), the thickness of the layer; ρ (Å) roughness of the layer; SLD (10^−6^ Å^−2^) scattering length density of the layer; Solvent % D_2_O penetration in each layer.

|  | d (Å) | ρ (Å) | SLD (10<sup>-6</sup> Å<sup>-2</sup>) | solvent % |
| --- | --- | --- | --- | --- |
| **Inner heads** | 5 ± 2 | 2 ± 1 | 1.87 | 5 ± 1 |
| **Lipid Chains** | 38 ± 3 | 1 ± 1 | -0.18 | 0.1 ± 0.1 |
| **Outer Heads** | 6 ± 2 | 2 ± 1 | 1.87 | 16 ± 4 |

While NR provides information on the average structure with respect to the bilayer normal, AFM can be used to resolve in detail the in-plane rafts morphology^37–40^. The SLB was formed on functionalized borosilicate glass coverslips, by injecting the liposomes (this time suspended in ultrapure water) in the buffer solution where they experienced an osmotic imbalance across the membrane, decreasing their pressurization (please refer to Materials and Methods for the details). As a result, following the adhesion to the substrate, liposomes will deform adopting more oblate shapes^41^, increasing the area occupied by each vesicle and favoring the previously described vesicle fusion mechanism. As shown in Figure 1c, consistently with NR data the surface is almost completely covered by a lipid bilayer, which presents nanometric domains of different heights, with the brighter areas corresponding to thicker membrane regions and the darker ones to thinner SLB portions. Accordingly, the height distribution of Figure 1d confirms the presence of two distinct lipid phases, with height values of h_d_ = 3.7 nm and h_o_ = 4.7 nm, in good agreement with the results obtained by Heberle et al. on the same vesicle preparation^15^. This thickness mismatch can be ascribed to the coexistence of two lipid phases of different composition, dictating variations in the membrane’s height^42–44^: in particular, membrane thickness was found to increase with length or degree of saturation of the lipid tails^43,44^. Here the thicker domains can be associated with the L_o_ phase, which is enriched with cholesterol and DSPC, i.e. a fully saturated long chain lipid. On the contrary, thinner regions correspond to the L_d_ lipid phase mainly composed of DOPC, which is characterized by a shorter tail length and two chain unsaturated bonds. After having properly flattened the image, by the application of a mask (see Figure S4), it is possible to determine the area fractions occupied by each of the two phases. Heberle et al.^15^ reported the area fraction corresponding to the L_d_ phase for liposomes of the very same composition to be 0.52; our calculations on SLBs are in line with those findings, giving a L_d_ area fraction of 0.50. Results also suggest that the SLB formation did not significantly modify the amount of L_d_ and L_o_ lipids, originally present in the unfused vesicles and that the lipid phase behavior is not affected by the presence of the solid support. The presented results corroborate the essential role of AFM in providing comprehensive morphological details on structure of rafted membranes. In the following paragraph, we extend the existing literature on AFM-based rafts characterizations^37–40^, by studying the structure of lipid rafts following their interaction with AuNPs.

### Interaction of AuNPs with lipid rafts: localization of AuNPs at the boundaries

In order to investigate the interaction of 16 nm citrated AuNPs (please refer to Materials and Methods for AuNP synthesis and to SI for AuNPs characterization details) with the lipid rafts present in the SLB, 5 μl of the NPs dispersion were injected in the ultrapure water buffer. Different literature reports connect the presence of phase segregation within the lipid bilayer to the selective adsorption of NPs along the domains boundaries^23,24^; however, a direct proof of this interaction is still missing to date. AFM represents one of the few techniques that could provide the sufficient resolution to simultaneously resolve the height difference between the two lipid phases (~ 1 nm) and the morphology of AuNPs. Despite the high resolution provided by AFM, the measurement remains challenging, as the spontaneous attachment of AuNPs to the probe (Figure S5) can often lead to imaging artifacts. In order to overcome this problem, the AFM fluid cell was filled with the same saline buffer used for SLB formation and the force SetPoint was kept on very low values (lower than ~ 200 pN). The use of the saline buffer as imaging solution should compensate the tip-sample electrical double-layer repulsion^32^ and limit the attachment of the NPs to the probe. In order to identify the portions of lipid bilayer characterized by the presence of AuNPs, images of 5×5 μm regions were initially acquired. Figure 2a shows a representative AFM topography of the SLB following the NPs injection. The bigger spherical objects represent vesicles that still have to fuse within the bilayer, while the smaller ones are the AuNPs, which seem to be homogeneously distributed above the SLB.

**Figure 2.**
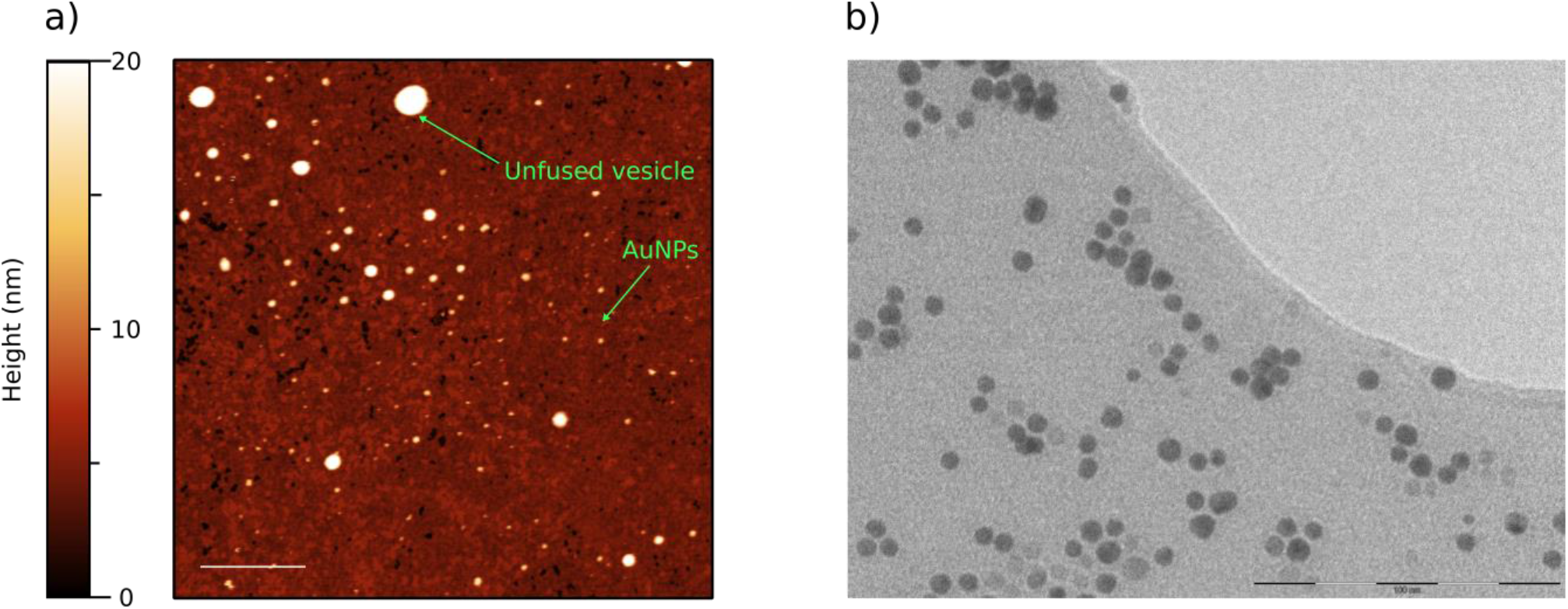
a) Representative AFM topography of the SLB following the injection of AuNPs. Lipid rafts are still visible as differently shaded areas. The larger and heterogeneous spherical objects represent unfused vesicles while the smaller ones are the AuNPs that have been homogeneously adsorbed on the lipid bilayer. Scalebar is 1 μm. b) Transmission Electron Microscopy (TEM) image of the AuNPs that were used in the experiments, scalebar is 100 nm (please refer to the SI for details regarding TEM characterization).

From a simple AFM topography, small lipid vesicles can be confused with AuNPs or AuNPs clusters; this could introduce statistical noise to the localization and morphometrical analysis. We recently developed an AFM-based nanomechanical characterization able to discriminate lipid vesicles from objects with the same morphology but different mechanical behavior^41^. This method evaluates the deformation that lipid vesicles undergo once adsorbed on a surface, by calculating their contact angle (α). Through the measurement of α and by assuming that the surface area of the vesicles is preserved upon adsorption, it is also possible to evaluate the diameter that characterizes the unperturbed vesicles in solution (called Diameter in solution). As described in Figure 3, lipid vesicles are characterized by a narrow distribution of contact angles over a wide range of sizes (Diameter in solution), while AuNPs present a narrow size distribution and higher contact angle values. This enables the easy singling out of the AuNPs and their exclusive inclusion in the next analysis.

**Figure 3.**
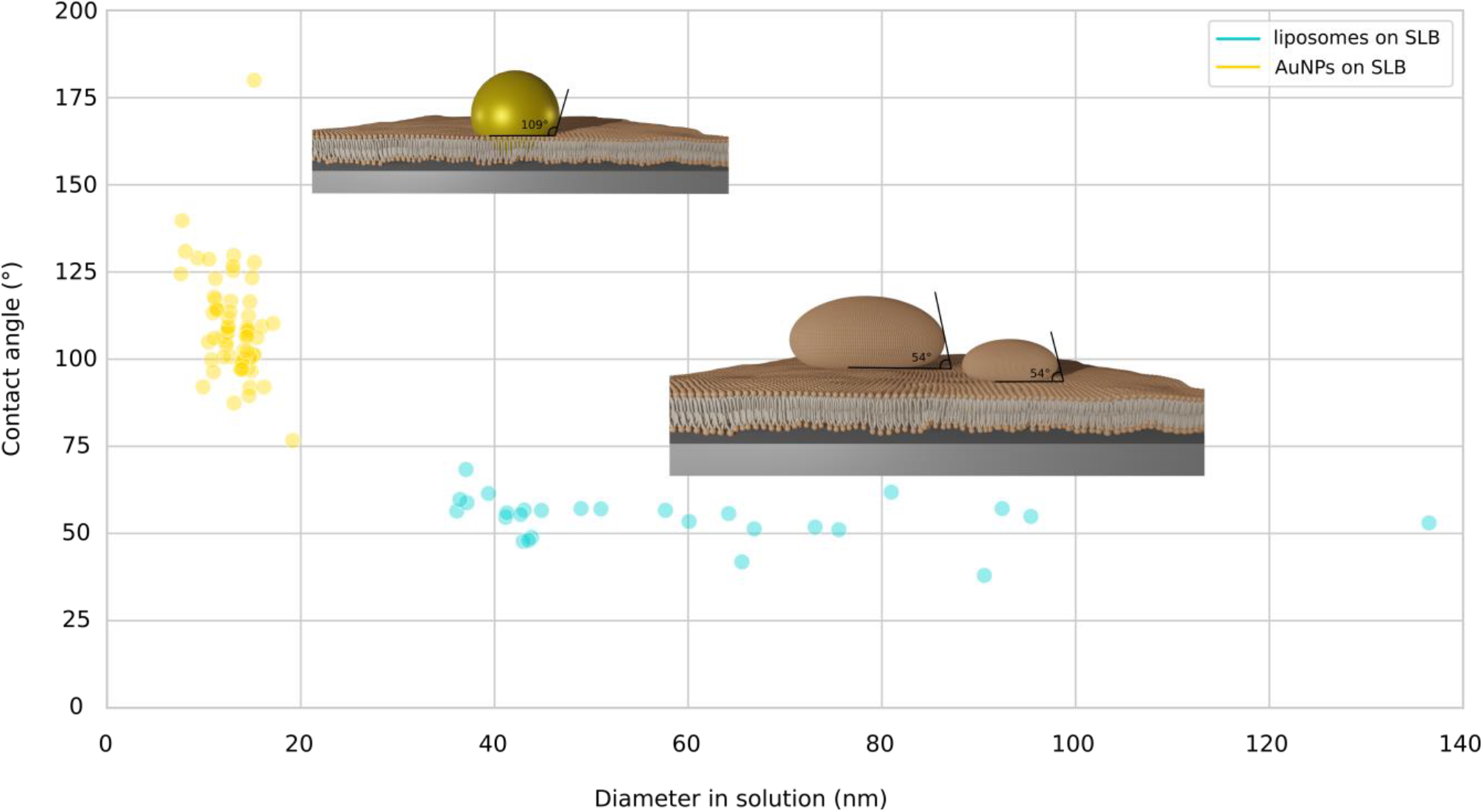
Plot representing the distributions of contact angle vs solution diameter of either vesicles (blue circles) and AuNPs (yellow circles). Vesicles data have been obtained from the liposomes present in Figure 2a while the AuNPs data come from micrographs like the ones reported in Figure 4a. Even though adsorbed on the SLB, liposomes show their nanomechanical fingerprint: a narrow contact angle distribution over a wide range of sizes. Their average contact angle is ~ 54° hence describing highly deformed shapes, possibly due to the SLB formation procedure. AuNPs display narrow distributions for both their size and contact angle, with average values of 14 nm and 109°, respectively.

In order to precisely determine whether the NPs targeted specific locations on the lipid matrix, the size of the scanned region was further reduced. In Figure 4a, representative images, with sizes of ~ 600×600 nm, illustrating the SLB decorated by AuNPs have been reported. The micrographs of Figure 4a constitute the direct proof of the AuNPs selective adsorption along the segregated phase boundaries.

**Figure 4.**
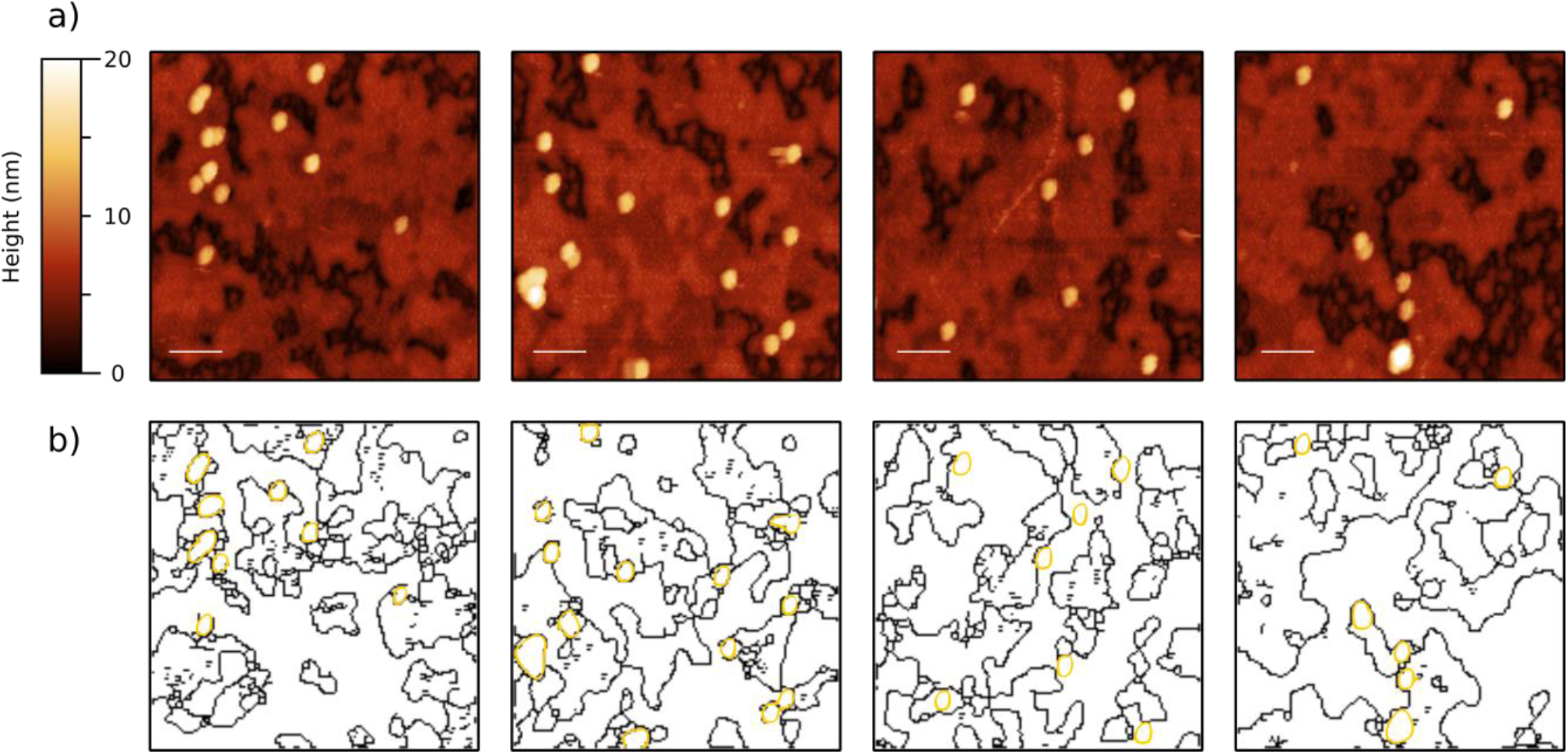
a) Representative AFM micrographs that clearly display the selective adsorption of AuNPs along the boundaries of the lipid rafts (brighter regions of the SLB that correspond to the L_o_ lipid phase). From the images it is also possible to distinguish between isolated and clustered NPs. All scalebars are 100 nm. b) Contour images obtained from the micrographs. Black lines represent the rafts edges while gold circles define the contours of the AuNPs. The gold NPs edges are always in contact with at least one of the lines describing the lipid segregated phase boundaries.

In the free image processing software Gwyddion 2.53.16, the sequential application of different masks allowed mapping the edges of either the lipid rafts and NPs shown in Figure 4a and, hence, obtaining a clearer indication of their relative positions. In Figure 4b the contour images of NPs and rafts have been superimposed with different colors, to highlight that AuNPs preferentially targeted the boundaries of the two lipid phases; indeed, the lines describing their shapes are always in contact with the edges of the lipid rafts. These results further corroborate the hypothesis that phase boundaries represent energetically favorable niches for lipid-NPs interactions. As previously discussed elsewhere^23^, NPs adsorption induces bilayer bending, which entails an energy penalty that increases the free energy associated with the overall process. This energy penalty is almost completely reduced along the phase boundaries, where the local negative curvature of the membrane, caused by the thickness mismatch between the two lipid phases minimizes the free energy associated with the NPs adsorption^23^.

### Inclusion of AuNPs within the lipid bilayer

AuNPs have a diameter of 16 nm (refer to SI for details), which is close to the average height measured with AFM imaging (14 ± 2 nm). This suggests that after adsorbing on the SLB, AuNPs probably penetrate the bilayer and reach the SiO_2_ surface. This result further extends the characterization of NPs-lipid interaction and corroborates our vision of rafts’ boundaries as regions of increased permeability^19–21^, where the membrane can easily wrap around the adsorbed NPs. Recent findings (Montis et al., JCIS, accepted on 31^st^ March 2020) confirm these results, suggesting that free-standing lipid bilayers can bend around the AuNPs surface, guided by citrate-lipid ligand exchange at the interface. All the above hypotheses are confirmed by the evaluation of the AuNPs contact angle with respect to the SLB. As suggested by Vinelli et al.^45^, the contact angle of a perfectly spherical, non-deformable (under the considered forces) object should be 180°, while we measured a substantially lower value. These apparent discrepancies can be rationalized by a careful morphological analysis, as detailed below.

The size of AuNPs is comparable with the tip radius, hence the effect of tip convolution should be taken into account. This was performed by assuming the NPs as perfectly spherical and non-deformable objects with heights that coincide with their actual diameters. This is a reasonable assumption given that, during an AFM measurement, the error along the vertical direction is negligible compared to the ones in the scanning plane. As a consequence, all the measured radii were then corrected by ~ 6 nm (corresponding to half the difference between the average NPs height and diameter measured by AFM). The NPs average contact angle, calculated with respect to the SLB and by using the corrected radii, gave a value of 109°, which is in very good agreement with the result that can be obtained from a simple geometrical model (Figure 5), featuring a 14 nm spherical and undeformable NP immersed in a ~ 5 nm lipid bilayer. For that case, α would be equal to 107°; this last result confirms that AuNPs penetrated the lipid bilayer and reached the underlying substrate.

**Figure 5.**
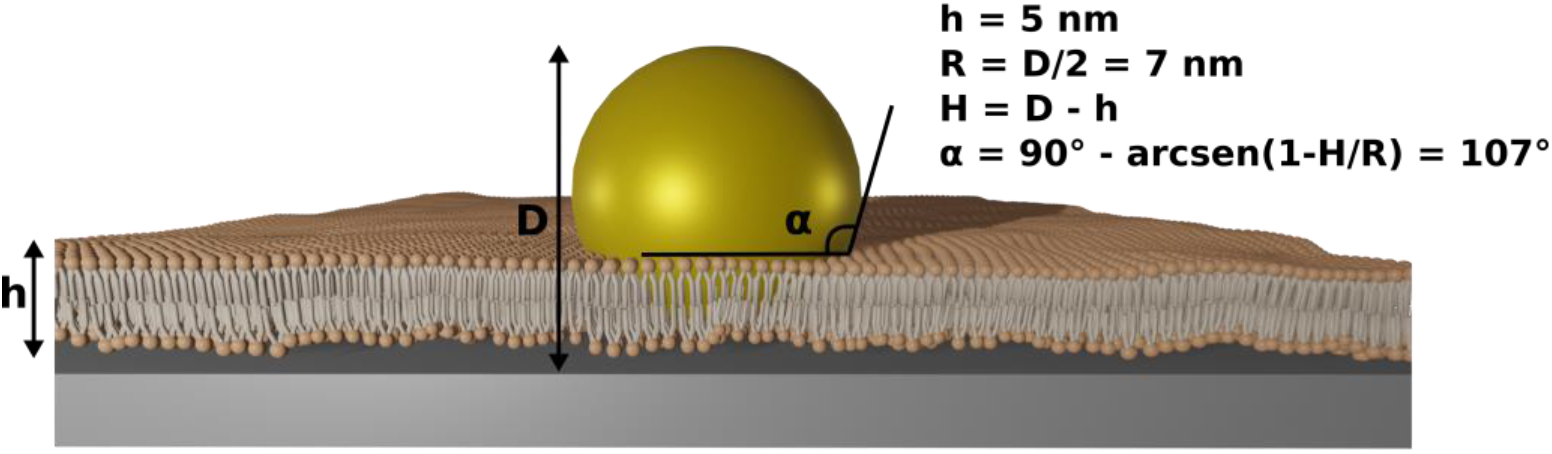
Schematic representation of the configuration used to evaluate, from a conceptual point of view, the contact angle that would characterize an AuNP with a diameter of 14 nm, adsorbed on a rigid flat surface and surrounded by a ~ 5 nm lipid bilayer.

## CONCLUSION

The presence of lipid rafts within the cell membrane has been linked with multiple important biological functions, like the formation and targeting of lipid nano-vesicles. The thickness mismatch that originates between the different immiscible segregated domains is thought to generate mechanical stresses, that enhance the membrane permeability along these regions. We herein exploited Atomic Force Microscopy to investigate the preferential adsorption of AuNPs along the phase boundaries of SLBs, generated from DOPC/DSPC/cholesterol (39/39/22 mol%) liposomes. Different works in the literature suggested a selective adsorption of AuNPs along the boundaries of lipid segregated domains, but a direct observation of this phenomenon is still missing to date. AFM allowed us to probe the existence of nanometric lipid rafts on the newly formed SLB and to spot the presence of NPs along their edges, hence providing a direct proof of this preferential adsorption pathway. In addition, we provided useful details about the experimental procedures that could significantly improve the reliability of AFM imaging; indeed, one of the major challenges hindering this type of measurements is the frequent tip contamination, caused by the attachment of the NPs to the AFM probe. We showed that the use of a saline buffer as imaging solution within the AFM fluid cell leads to optimal image quality and strongly reduces tip contamination events. Then, through the application of an AFM-based morphometric nanomechanical characterization, it was also possible to further investigate the reorganization of the lipid bilayer, as a consequence of the AuNPs adsorption. We found out that the lipid matrix wrapped around the NPs, allowing them to penetrate within the hydrophobic region until reaching the rigid SiO_2_ surface of the slides. The theoretical calculation of the morphological parameters describing this phenomenon is in perfect agreement with the experimental results and further corroborates our interpretation. Further studies will focus on extending this characterization to membranes with varying compositions and employing NPs of different core and/or size.

## AUTHOR INFORMATION

### Author Contributions

The manuscript was written through contributions of all authors. All authors have given approval to the final version of the manuscript.

### Notes

The authors declare no competing financial interests.

## ACKNOWLEDGMENTS

This work was also supported by the Consorzio Sistemi a Grande Interfase (CSGI) through the evFOUNDRY project, Horizon 2020-Future and emerging technologies (H2020-FETOPEN), ID: 801367. We thank the SPM@ISMN research facility for support in the AFM experiments. Maier-Leibnitz Zentrum is acknowledged for provision of beam-time.

## SUPPLEMENTARY INFORMATION

### AuNPs characterization

#### 1. TRANSMISSION ELECTRON MICROSCOPY

Trasmission electron microscopy (TEM) images were acquired with a STEM CM12 Philips electron microscope equipped with an OLYMPUS Megaview G2 camera, at CeME (CNR Florence Research Area, Via Madonna del Piano, 10 - 50019 Sesto Fiorentino). Drops of citrated AuNP, diluted ten times, were placed on 200 mesh carbon-coated copper grids with a diameter of 3 mm and a thickness of 50 μm (Agar Scientific) and dried at room temperature. Then, samples were analyzed at an accelerating voltage of 100 keV.

#### 2. SMALL ANGLE X-RAY SCATTERING (SAXS)

SAXS measurements were carried out on a S3-MICRO SAXS/WAXS instrument (HECUS GmbH, Graz, Austria) which consists of a GeniX microfocus X-ray sealed Cu Kα source (Xenocs, Grenoble, France) of 50 W power which provides a detector focused X-ray beam with λ = 0.1542 nm Cu Kα line. The instrument is equipped with two one-dimensional (1D) position sensitive detectors (HECUS 1D-PSD-50 M system), each detector is 50 mm long (spatial resolution 54 μm/channel, 1024 channels) and cover the SAXS q-range (0.003< q <0.6 ^−1^). The temperature was controlled by means of a Peltier TCCS-3 Hecus. The analysis of SAXS curves was carried out using Igor Pro^46^. SAXS measurements on AuNP aqueous dispersions, was carried out in sealed glass capillaries of 1.5 mm diameter. To analyze AuNPs profiles, we chose a model function with a spherical form factor and a Schulz size distribution^47^, which calculates the scattering for a polydisperse population of spheres with uniform scattering length density. The distribution of radii (Schulz distribution) is given by the following equation:

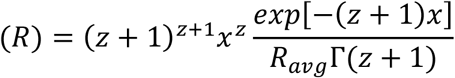

where R_avg_ is the mean radius, x = R/R_avg_ and z is related to the polydispersity of the dispersion. The form factor is normalized by the average particle volume, using the 3^rd^ moment of R:

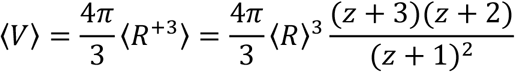

The scattering intensity is:

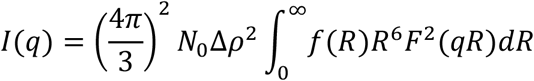

where N_0_ is the total number of particles per unit volume, F(R) is the scattering amplitude for a sphere and Δρ is the difference in the scattering length density between the AuNP and the solvent.

The structural parameters of citrated gold nanoparticles were evaluated from the SAXS profile of Figure S1 according to the above model.

**Figure S1.**
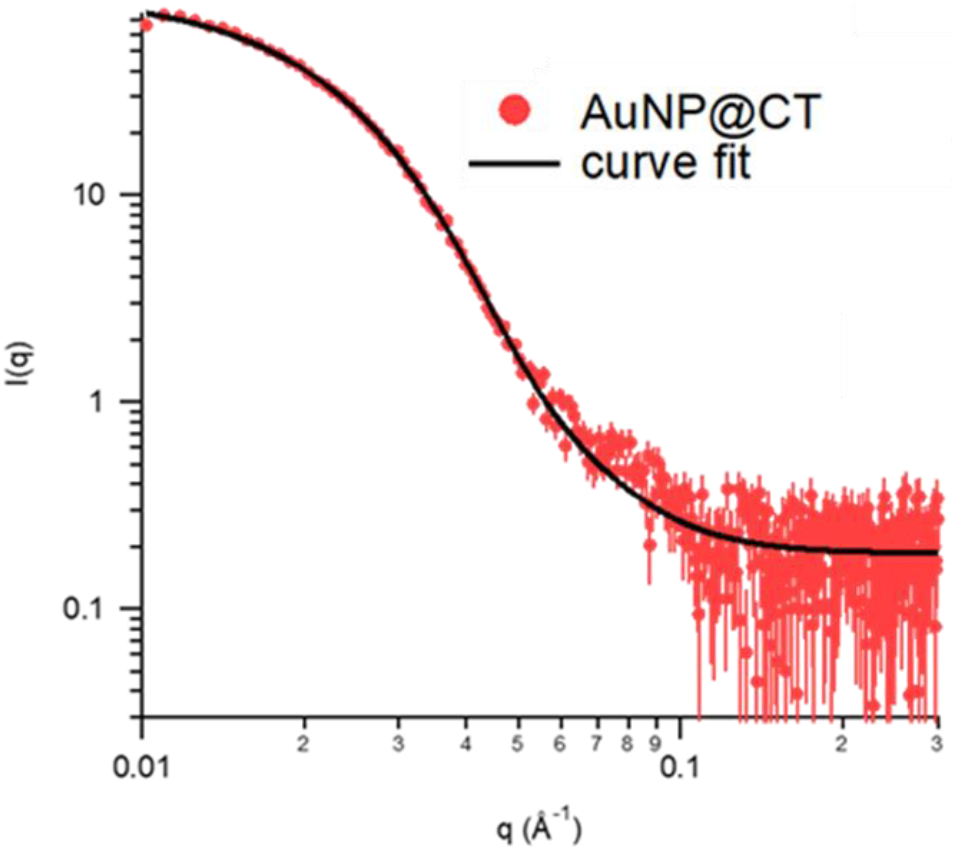
Experimental SAXS profile (markers) obtained for citrated AuNPs and curve fit (solid black line) according to the Schulz spheres model from the NIST package SANS Utilities. The size and polydispersity obtained from the fitting procedure are 13 nm and 0.3, respectively.

#### 3. UV-VIS SPECTROSCOPY

UV-Vis spectra were measured with a JASCO UV-Vis spectrophotometer.

The size of citrate gold nanoparticles was further evaluated from UV-Vis Spectroscopy by the following equation^48^:

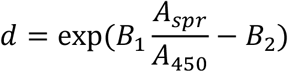

with d diameter of gold nanoparticles, A_spr_ absorbance at the surface plasmon resonance peak, A_450_ absorbance at the wavelength of 450 nm and B_1_ and B_2_ dimensionless parameters, taken as 3 and 2.2, respectively. The obtained diameter value is 16 nm.

**Figure S2.**
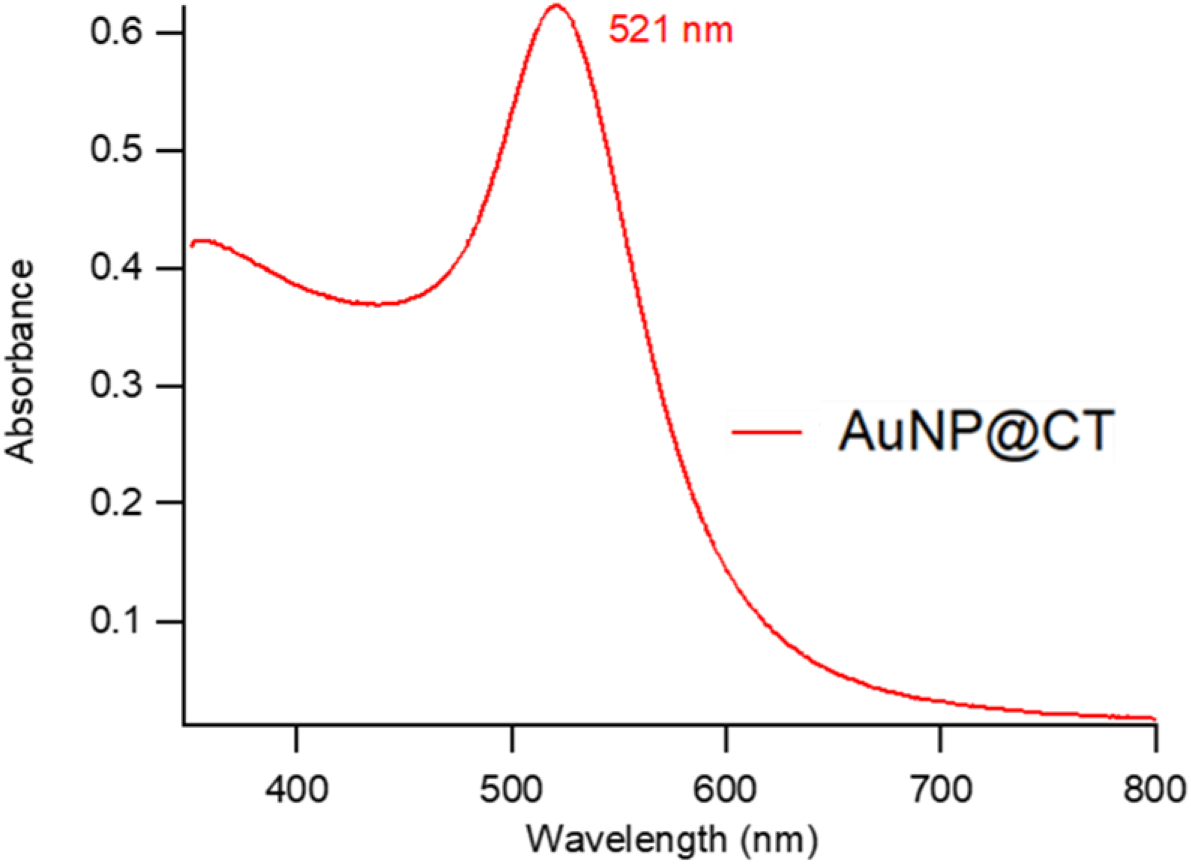
UV-Vis absorption spectra of citrated AuNP dispersion (after 1:5 dilution in water). The plasmon absorption peak is located at 521 nm.

The concentration of citrated gold nanoparticles was determined via UV-Vis spectrometry, using the Lambert-Beer law (E(λ) = ε(λ)lc) and considering the extinction values *ε* (λ) at the LSPR maximum, i.e. λ = 521 nm. The extinction coefficient ε(λ) was determined by the following equation^49^:

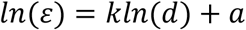

with d core diameter of nanoparticles, and k and a dimensionless parameter (k = 3.32111 and a = 10.80505). The arithmetic mean of the size, obtained by both the optical and the scattering analyses, leads to an ε(λ) value of 4.8·10^8^ M^−1^cm^−1^. Consequently, the final concentration of citrated AuNP is ~7.8·10^−9^ M.

#### 4. ATOMIC FORCE MICROSCOPY

Preliminary AFM images of AuNPs adsorbed on bare mica substrates were performed in order to test the effect of using a saline buffer as imaging solution, to prevent the attachment of NPs to the AFM tip (please refer to the Materials and Methods section for details regarding the imaging setup and parameters).

**Figure S3.**
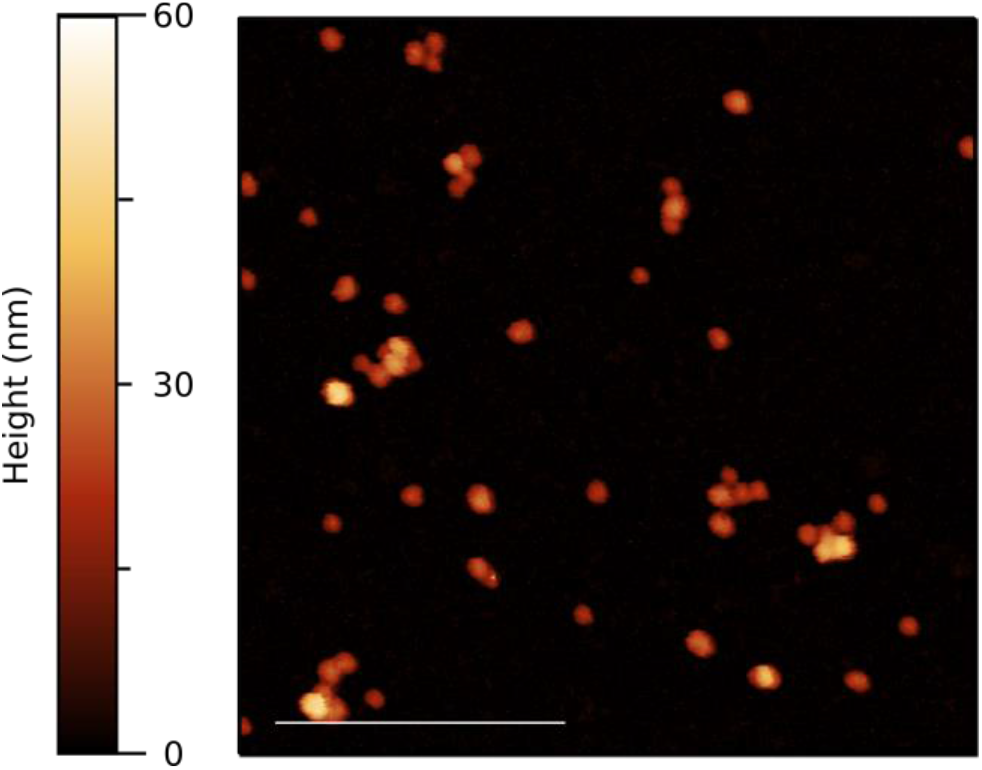
AFM topography representing AuNPs either isolated or clustered, adsorbed on a bare mica substrate. As reported in the Materials and Methods section, AFM imaging was performed using a saline buffer as imaging solution, in order to obtain better image quality and avoid tip contamination due to the attachment of the NPs to the probe. As can be seen from the image, isolated AuNPs get faithfully described by nearly perfect spherical shapes with an average contact angle close to 180°. Scalebar is 400 nm.

### Neutron Reflectivity measurements

The software MOTOFIT was employed for the analysis of the NR curves. A five-layer model was employed to analyze the reflectivity profiles of neat SLBs, with scattering length density values calculated for each layer: a bulk subphase of Si (SLD = 2.07 * 10^−6^ Å^−2^), a superficial layer of SiO_2_ (SLD = 3.47 * 10^−6^ Å^−2^); a second layer of D_2_O (SLD = 6.393 * 10^−6^ Å^−2^); a third layer composed of the polar headgroups of the SLB of the inner leaflet (SLD = 1.87 * 10^−6^ Å^− 2^); a fourth layer composed of the bilayer’s lipid chains (SLD = −0.18 * 10^−6^ Å^−2^); a fifth layer composed of the polar headgroups of the outer bilayer’s leaflet (SLD = 1.87 * 10^−6^ Å^−2^); a bulk superphase of solvent (D_2_O, SLD = 6.393 * 10^−6^ Å^−2^). The scattering length density values for the polar headgroups and lipid chains were estimated by taking into account the chemical compositions and the submolecular fragment volumes of phosphatidylcholines as determined by Armen et al. through molecular dynamic simulations^50^ (see table S1).

**Table S1.** Chemical formula, molecular volumes and corresponding scattering length densities of species relevant to this study.

| **Molecule** | **Chem. Formula** | **Volume (Å<sup>3</sup>)** | **SLD(10<sup>-6</sup> Å<sup>-2</sup>)** |
| --- | --- | --- | --- |
| **PC headgroups** | C<sub>10</sub>H<sub>18</sub>NO<sub>8</sub>P | 321.9 | 1.866 |
| **DSPC chains** | C<sub>34</sub>H<sub>70</sub> | 1004.6 | -0.357 |
| **DOPC chains** | C<sub>34</sub>H<sub>66</sub> | 982.2 | -0.212 |
| **cholesterol** | C<sub>27</sub>H<sub>46</sub>O | 630 | 0.210 |

**Figure S4.**
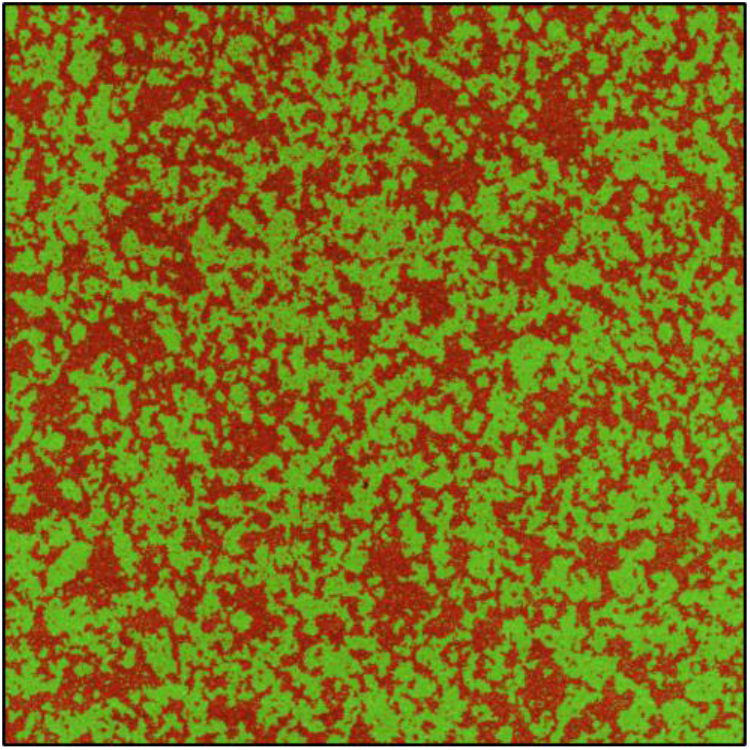
AFM topography of the SLB obtained from DOPC/DSPC/Chol (39/39/22 %w/w) liposomes. Using Gwyddion 2.53.16 it was possible to apply a mask to selectively cover the Lo phase (characterized by higher height values) and estimate the area fraction of each phase. The area fractions of the L_o_ and L_d_ phases are approximately 0.50, confirming the results obtained by Heberle et al.^15^ on the very same vesicles preparation.

**Figure S5.**
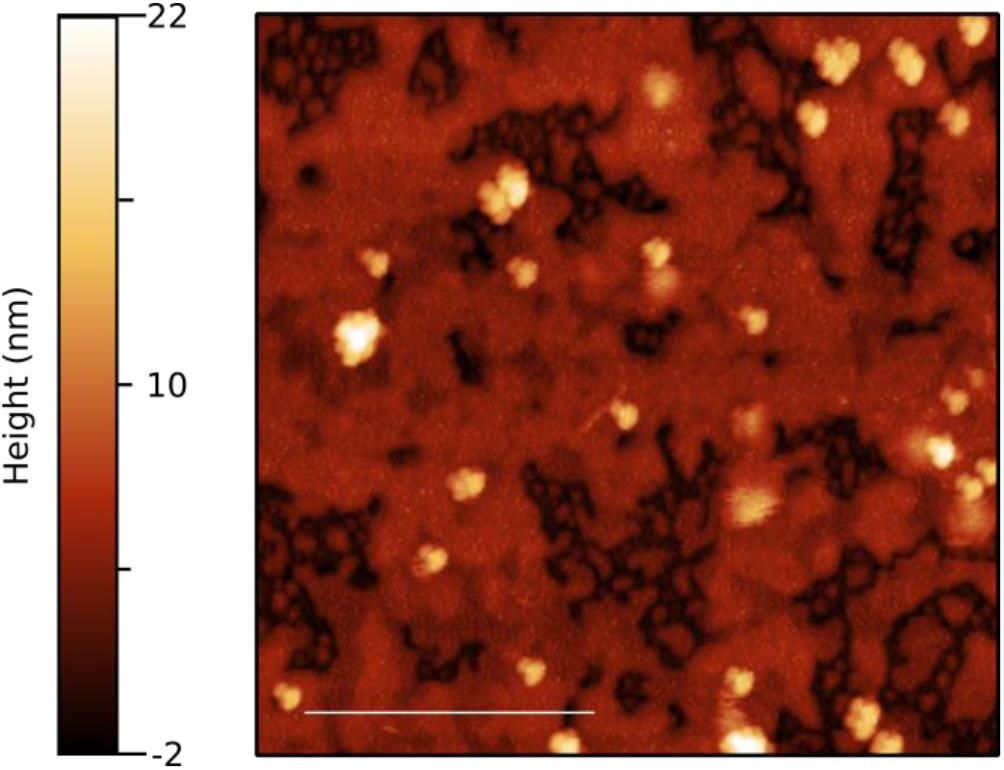
AFM topography showing the effects of tip contamination on the imaging of AuNPs adsorbed on the lipid bilayer. While the SLB gets correctly imaged, all the NPs appear in “clusters” characterized by similar shapes (same protrusions in all the directions). This is a clear indicator that the AFM probe has been contaminated by the attachment of one or multiple NPs which lead to the generation of imaging artifacts. Scalebar is 400 nm.

